# DNA-binding factor footprints and enhancer RNAs identify functional non-coding genetic variants

**DOI:** 10.1101/2023.11.20.567860

**Authors:** Simon C Biddie, Giovanna Weykopf, Elizabeth F. Hird, Elias T. Friman, Wendy A Bickmore

**Affiliations:** MRC Human Genetics Unit, Institute of Genetics and Cancer, University of Edinburgh, UK; NHS Lothian, Edinburgh, UK

## Abstract

Genome-wide association studies (GWAS) have revealed a multitude of candidate genetic variants affecting the risk of developing complex traits and diseases. However, these highlighted regions are typically in the non-coding genome, and uncovering the functional causative single nucleotide variants (SNVs) is challenging. Prioritisation of variants is commonly based on functional genomic annotation with markers of active regulatory elements, but current approaches still poorly predict functional variants. To address this, we systematically analyse six markers of active regulatory elements for their ability to identify functional variants. We benchmark against molecular quantitative trait loci (molQTL) from assays of regulatory element activity that identify allelic effects on DNA-binding factor occupancy, reporter assay expression, and chromatin accessibility. We identify the combination of DNase footprints and divergent enhancer RNA as markers for functional variants. This signature provides high precision, trading-off low recall, thus substantially reducing candidate variant sets to prioritise variants for functional validation. We present this as a framework called FINDER – Functional SNV IdeNtification using DNase footprints and Enhancer RNA, and demonstrate its utility to prioritise variants using leukocyte count trait and analyse variants in linkage disequilibrium with a lead variant to predict a functional variant in asthma. Our findings have implications for prioritising variants from GWAS, in development of predictive scoring algorithms, and for functionally informed fine mapping approaches.

## Introduction

Only a very small (2%) proportion of the human genome codes for protein-coding genes. Yet, genetic variants in the non-coding genome are known to contribute to Mendelian diseases and complex traits. Genome-wide association studies (GWAS) have uncovered thousands of genetic variants that are associated with traits or alter disease risk, and these loci predominantly lie in the non-coding genome (Maurano et al., 2012). Identifying functional and causal non-coding variants from GWAS, and the target genes they regulate, remains a major challenge. The model for effects on trait proposes that non-coding variants alter the target sequences of sequence-specific nucleic acid binding proteins in regulatory elements resulting in quantitative changes in target gene expression. This could be at the level of modulating RNA splicing by RNA binding proteins, (Qi et al., 2022), or through altering the affinity for DNA binding proteins such as transcription factors (TF) at enhancers (Maurano et al., 2015; Johnston et al., 2019). Rare, pathogenic mutations in enhancers have been shown to reduce (Jeong et al., 2008) or increase (Lettice et al., 2012) TF binding. Contrasting the large functional affects for rare alleles, common single nucleotide variants (SNVs) tagged in GWAS are thought to modulate TF occupancy at regulatory elements (Kasowoski et al., 2010; Maurano et al., 2015; Carrasco Pro et al, 2020).

Because GWAS identifies regions associated with disease risk that contain multiple SNVs in linkage disequilibrium (LD), identifying the functionally relevant SNV(s) poses a significant challenge. One route is to analyse changes in chromatin structure at active regulatory elements. Binding of TFs at enhancers leads to nucleosome displacement, and recruitment of co-activators that can then further modify chromatin including, but not limited to, acetylation of histone H3 lysine 27 (H3K27ac) (Vernimmen and Bickmore, 2015). The Assay for Transposase-Accessible Chromatin using sequencing (ATAC-seq) is widely used for profiling sites with disrupted nucleosome structure and has been used in efforts to prioritise risk variants (Grishin and Gusev, 2022; Su et al., 2022). Prior to the widespread use of ATAC-seq, DNase hypersensitive site-seq (DHS-seq) and DNase I footprinting have been used to identify sites of likely TF-binding and nucleosome disruption and to assess the effect of genetic variants on TF binding (Maurano et al., 2015; Moyerbrailean et al., 2016; Vierstra et al., 2020).

Enhancers and their target genes are most often located in the same topologically associating domain (TAD). Therefore, the integration of GWAS with data on tissue-specific features of active chromatin (ATAC-seq, DHS-seq, DNase footprinting, H3K27ac ChIP-seq), and chromosome conformation capture, has been used to try and predict functional variants and their target genes (Schaub et al., 2012; Nasser et al., 2021; Gazal et al., 2022). However, predictions based on active regulatory marks often fall short in pinpointing functional variants (Gasperini et al., 2017, Chen et al., 2023)

Computational models to predict functional non-coding variants using machine learning methods including deep neural networks or linear regression models based on feature annotations, such as open chromatin, histone marks, TF binding, and conservation scores have also been developed (Wang et al., 2022). However, whilst showing some efficacy for rare variants, they poorly predict non-coding variant effects for common variants (Wang et al., 2022, Tabarini et al., 2022). Prioritising and identifying functional non-coding variants therefore remain a barrier to GWAS follow-up studies.

Another feature of active enhancers is the production of short, unstable bidirectional enhancer RNAs (eRNAs) (Hou and Krause, 2021). To the best of our knowledge this feature has not been used in the prediction of functional genetic variants. Here, we analyse the utility of active regulatory element marks – open chromatin, H3K27ac, DNase footprints, TF binding, as well as the presence of divergent enhancer RNA (eRNA), for the prioritisation and discovery of functional variants. We analysed active regulatory markers using > 50,000 datasets from > 2,000 cell types, tissue types, and treatment conditions and benchmarked these features with variants that demonstrate enhancer modulating activity using *cis*-acting molecular quantitative trait loci (molQTL). We find that the combination of DNase footprints and eRNA has high precision for predicting functional non-coding variants, and we apply this approach to data from GWAS traits. In this way we demonstrate a framework to allow prioritisation of candidate variants from GWAS data to aid discovery of functional and causative genetic variants.

## Results

### A compendium of genomic markers of active regulatory regions

Functional non-coding genetic variants are thought to reside in regulatory elements in the non-coding genome. Based on this, we collated a compendium of published datasets of markers for active regulatory elements from human cells and tissues. We included markers for open chromatin (DNase hypersensitive sites (DHS) from DNase-seq, and Assay for Transposase-Accessible Chromatin (ATAC-seq)), H3K27ac, divergent enhancer RNA (eRNA), DNase footprints, and DNA-binding factor occupancy from chromatin immunoprecipitation (ChIP). Data were collected from uniformly processed resources, which included multiple replicates, across cell lines, primary cells or tissue biosources (In aggregate this comprised > 50,000 datasets from > 2,000 biosources (Figure 1A, Supplementary Table 1). Most datasets are from ChIP-seq, including ChIP for 956 DNA-binding factors and for some factors across multiple cell types or treatment conditions. Only one cell-type, hepatocytes, shares data across all types of assays (Supplemental Figure 1A).

**Figure 1.**
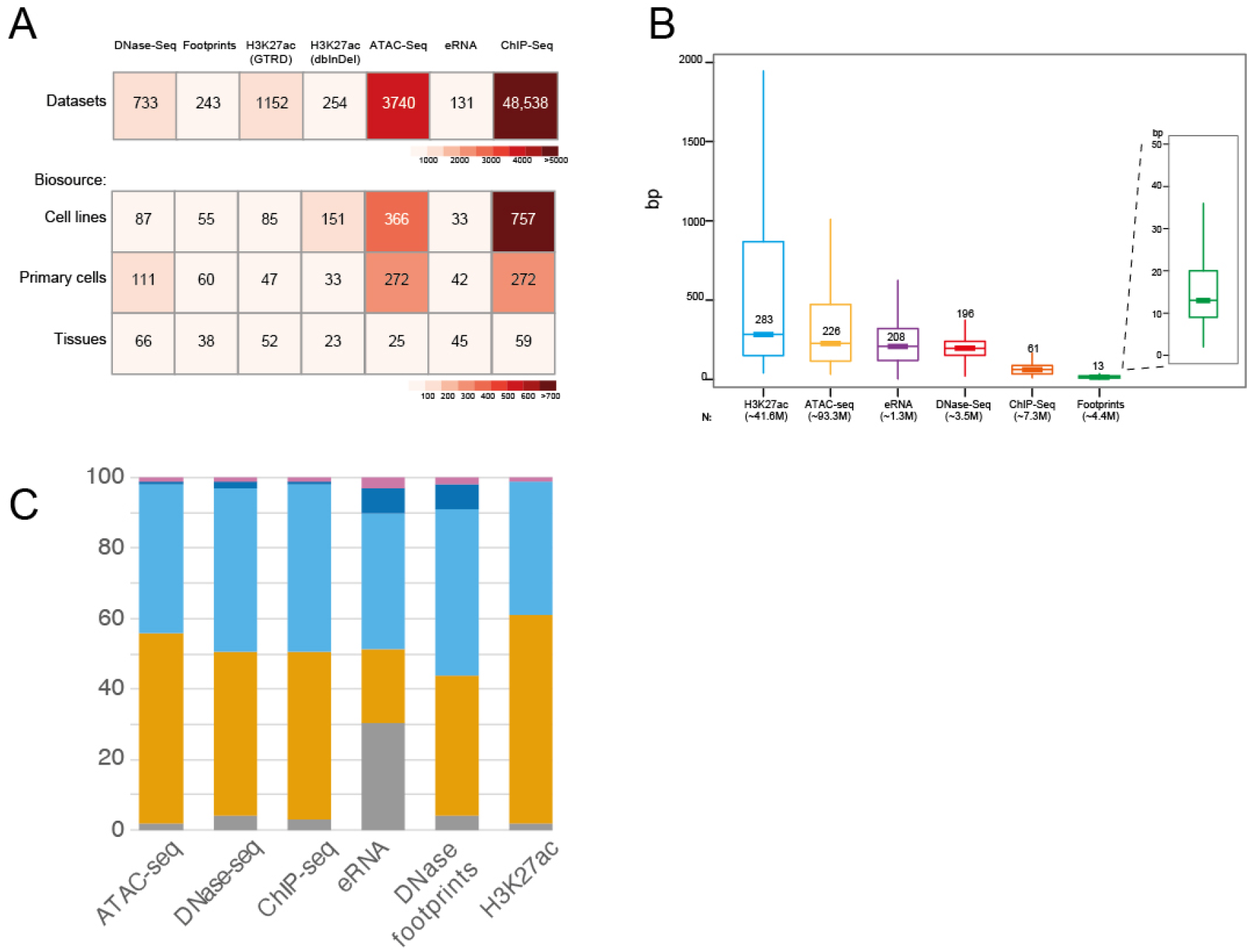
A compendium of markers of active regulatory regions. A. Heatpmap for six datasets that mark active regulatory elements. H3K27ac data were obtained from two different sources. Footprints are derived from DNase-seq data. The number of biosources (cell lines, primary cells, or tissues) are indicated. For ChIP-seq datasets, this consists of 956 DNA-binding factors. B. Boxplot for each feature length from each biosource with the number shown indicating the median length in basepairs. The total number of elements for each feature is shown, as a sum for all biosources regardless of overlapping features. C. Genomic annotations for markers of active regulatory markers from merged biosources.

Taking the aggregate across all biosources for each marker, the number of elements ranges from 1.3 million (eRNA), to 93.3 million (ATAC-seq), with the median length of the elements dependent on the assay (Figure 1B). The majority of active regulatory elements are found in the non-coding regions - intergenic regions, promoters or introns (Figure 1C).

For each marker, we merged overlapping datasets across all biosources to determine the non-redundant genome coverage of active regulatory elements. The largest coverage is represented by ATAC-seq (75.2%), followed by H3K27ac (61.5%). This is likely due to the high number of biosources (Figure 1A), the longer median length and distribution (Figure 1B), but also probably indicates too low a threshold used in peak calling. The smallest coverage is from eRNA (2.1%) followed by DNase footprints (3.6%) (Supplemental Figure 1B).

### Prioritisation of genetic variants by combinations of regulatory markers

Functional non-coding genetic variants are likely to impact target gene expression through altering activity at regulatory elements. To assess the utility of markers of active regulatory elements in prioritising functional non-coding variants, we established a set of variants which colocalise with molecular quantitative trait loci (molQTL) to benchmark against. We selected three molQTL assays which measure different molecular mechanisms of regulatory element activity and which demonstrate variant effects in *cis* that impact regulatory element activity – DNA-binding factor QTL (bQTL) (Abramov et al., 2021), massively parallel reporter assay (MPRA) QTL (raQTL) (van Arensbergen et al., 2019), and chromatin accessibility QTL (caQTL) (Vierstra et al., 2020, Zheng et al., 2020) (Figure 2A). Variants identified by these molQTL assays are derived from >16K datasets across >800 biosources (Figure 2B, Supplementary table 1). The majority of these datasets are from bQTL assays based on ChIP-seq annotations, while caQTL are based on DNase-seq and ATAC-seq datasets. raQTL variants were obtained from one resource which identified raQTLs in K562 and HepG2 cell lines, using an unbiased set of 5.9 million variants (van Arensbergen et al., 2019). We have not included other studies using MPRA to identify raQTL as these often use variants identified from specific GWAS traits which would bias our downstream analysis. There was minimal overlap between variants colocalising with bQTL, raQTL and caQTL (Figure 2C). This maintains each molQTL assay as a largely independent set of variants to benchmark for functional variants.

**Figure 2.**
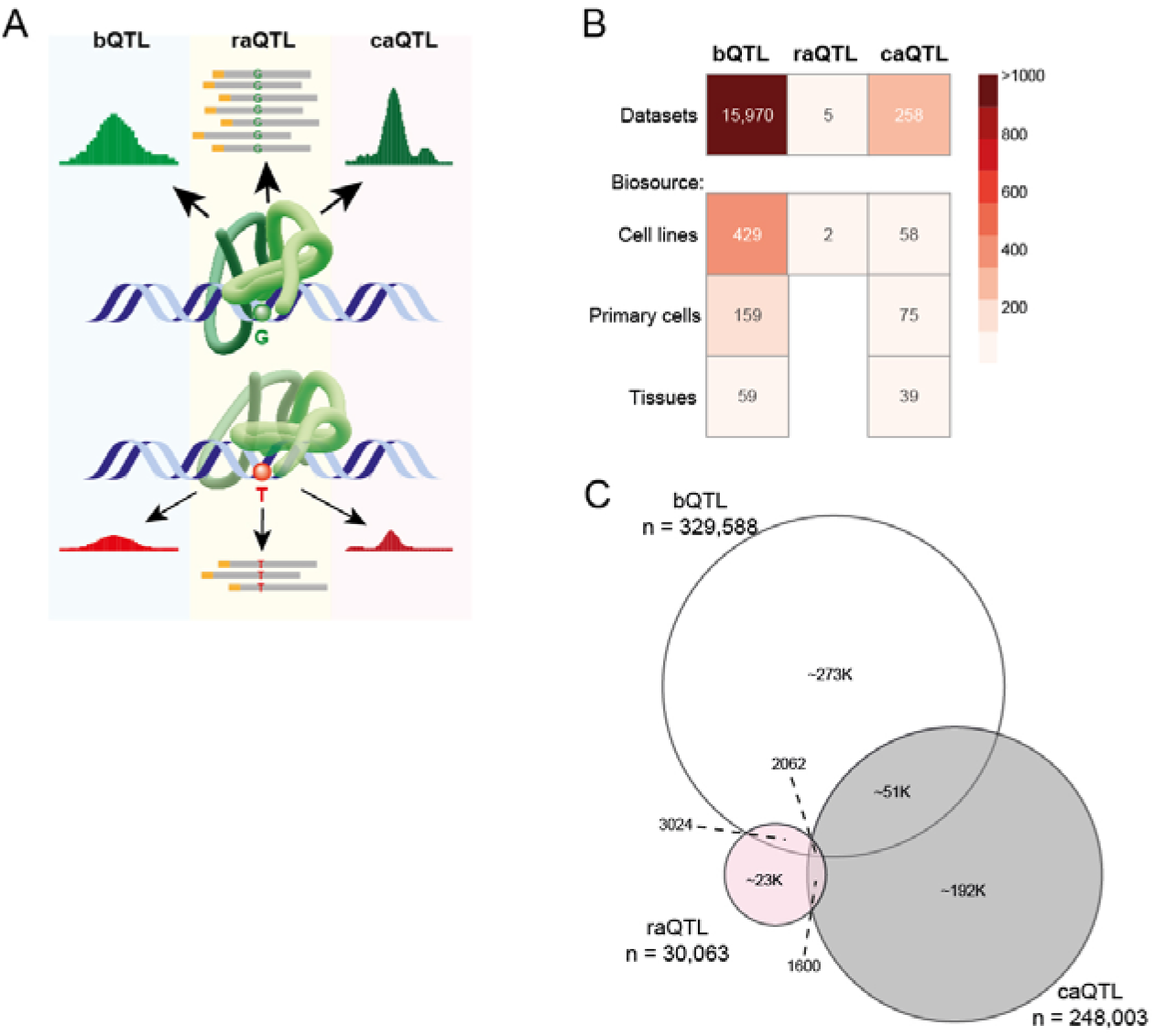
Molecular quantitative trait loci (molQTL) with effects on regulatory element activity. A. MolQTL with effects on regulatory element activity include binding QTLs (bQTL), reporter assay QTLs (raQTL) from massively parallelel reporter assays (MPRA), and chromatin accessibility QTLs (caQTL). The model depicts binding of a DNA-binding factor such as a transcription factor (TF), which is altered by a single nucleotide variation (SNV), thus reducing binding, reporter assay expression and chromatin accessibility. B. Heatmap showing the molQTL datasets and biosources. bQTL data are derived from ChIP-seq experiments of 1073 DNA-binding factors. C. Proportional Venn diagram of the number of QTLs from each assay showing minimal overlap.

Using each molQTL to benchmark, we aimed to identify functional variants from the GWAS catalogue (Sollis et al., 2023), which is derived from >28K studies and >5K traits including diseases, containing variant-trait associations with p-value of ≤ 1.0 × 10^-5^ (Supplementary table 1). This set contains >180K non-redundant variants, the majority of which are in non-coding regions (Supplemental Figure S2A). To identify how active regulatory marks can functionally annotate non-coding variants, we merged all biosources for each mark, and in the case of ChIP this included merging all DNA-binding factors (Supplemental Figure 2B). This overcomes two main issues. Firstly, that the cell- or tissue-type in which a variant is functional is not known *a priori*. Secondly, the low overlap of biosources for each marker (Figure 1B) would limit intersection analysis. We therefore took a cell/tissue agnostic approach by merging biosources. With this, we determined the co-localisation of the GWAS catalogue variants with each marker of active regulatory elements, and all combinations of markers, totalling 55 combinations from two-way up to six-way combinations.

We determined the precision (positive predictive value (PPV) – [true positive / (true + false positives)]) for each marker and their combinations by the intersection with variants from the GWAS catalogue, benchmarked against the intersection of the marker combinations with each molQTL as a true positive set. The precision value was dependent on the size of the molQTL set relative to the larger GWAS catalogue. The range of precision values were 7.5 – 55.46% for bQTL, raQTL;0.42 - 3.53%, and caQTL; 4.02 – 27.7%. In all cases the highest precision was achieved by the combination of all six markers. To compare precision values between the molQTL, we converted precision values to Z-scores for each molQTL, giving a range of scores from -2.2 to 1.44. Using unsupervised clustering we generated a Z-score heatmap of precision values for feature combinations against molQTL assays, showing four clusters (Figure 3A). The cluster with the highest Z-scores (cluster 3) showed that two markers, DNase footprints and eRNA, were present for all feature combinations within the cluster, including the combination of these two markers alone. In contrast, the cluster with the lowest Z-score (cluster 1) excluded any combination of markers that had footprints or eRNA. As independent markers, DNase footprint or eRNA, were present in an intermediate cluster (cluster 2). This suggests that the combination of footprints and eRNA, with or without other markers, provides high precision for identifying functional variants. Increasing the number of features increased the number of predicted negative variants, although the number of predicted negative variants is >90% for footprints or eRNA alone (Supplemental Figure 3).

**Figure 3.**
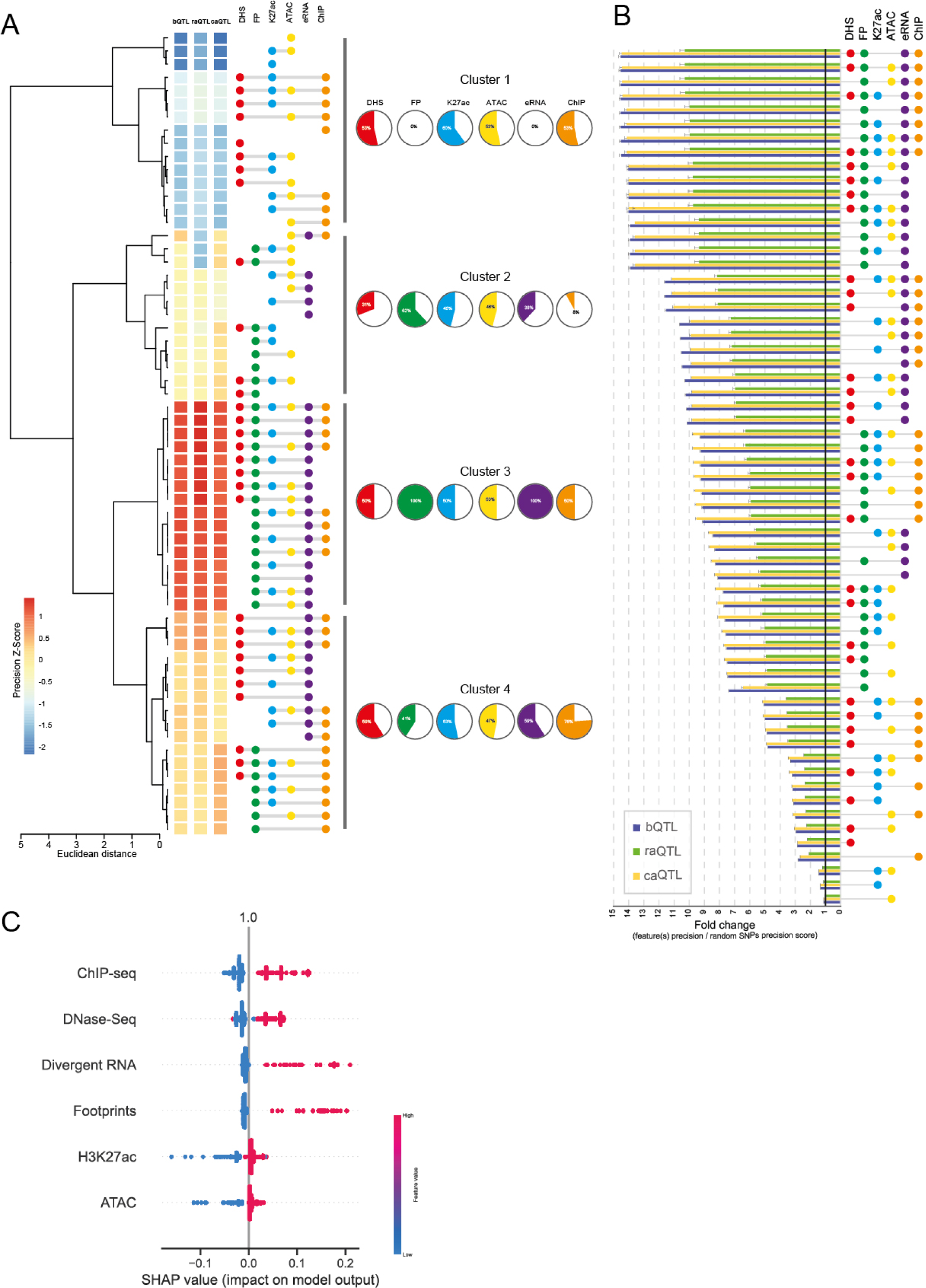
Identifying functional variants using combinations of markers of active regulatory elements. A. Heatmap of precision Z-scores for individual, or combinations of, markers for GWAS catalogue variants. Variants from the GWAS catalogue were intersected with each feature or combinations of features, benchmarked against molQTLs. Unsupervised clustering was performed and percentages of each feature represented in each cluster is shown. B. Fold change of precision scores for functional variant discovery by feature combinations from random genomic SNVs. Common (MAF >= 1%) SNVs from the whole genome were subsetted to a comparable number to the GWAS catalogue. Precision scores for variants that intersect with feature(s) and co-localise with a molQTL, are expression as fold change relative to the number of subsetted variants that co-localise with molQTL. Feature combinations are sorted from lowest to highest fold change. 100 iterations of subsetting from dbSNP were performed and error bars shows standard deviation. C. SHAP (SHapley Additive exPlanations) values for a machine learning model using a random forest classifier to identify feature importance for each active regulatory mark towards the functional variant predictive model.

We next considered the precision of marker combinations to identify functional variants across the entire genome, without association with GWAS traits. Taking all common genetic variants (minor allele frequency (MAF) >= 1%), consisting of >660M non-redundant single nucleotide variants (SNV), we randomly selected subsets to obtain a comparable number to the GWAS catalogue of ∼180K variants. We calculated the precision of combinations of markers and then converted this to a fold change over random SNVs. We repeated the random SNV subsetting 100 times to generate a mean fold change and plotted the markers and combinations from lowest to highest fold change (Figure 3B). This showed that ATAC-seq, H3K27ac, and the combination of the two, are poor discriminators of functional variants (mean fold change range; 1.12 to 1.44). In comparison, a step change is observed for the combination of DNase footprints and eRNA, with a mean fold change of 13.89 over random SNVs. Footprints and eRNA alone show a mean fold change of 7.37 and 8.09 respectively, suggesting that they can individually provide functional annotation of variants across the genome, but with higher precision when combined. This is in keeping with the precision observed from footprints with eRNA to identify functional variants in the GWAS catalogue. Our analyses suggest that the combination of footprints and eRNA are strong predictors of functional variants with high precision, with additional markers providing only small increments in precision. Importantly, when considering the precision of GWAS compared to random genomic variants, the GWAS catalogue shows a mean fold change of 8.24 which, perhaps expectedly, demonstrates that GWAS can identify functional variants, above a random model.

To corroborate our findings for footprints and divergent RNA in predicting functional variants, we employed an independent machine learning approach. We trained a Random Forest Classifier model to predict any molQTL signature (bQTL/raQTL/caQTL) based on overlaps with the six active regulatory markers, providing a predictive value of 0.64. We determined SHAP values as a measure of feature importance for the model, which revealed that footprints and divergent RNA are the top positively associated features for the predictive model (Figure 3C).

### Centrality in open chromatin to predict functional non-coding variants

TF binding initiates and maintain chromatin accessibility (Biddie et al., 2011) and indeed summits of DHS have been shown to be enriched for DNA-binding factor motifs (Vierstra et al., 2020). Using a consensus reference map of DHS with summit annotations (Meuleman et al., 2020), we tested if centrality in DHS could predict functional variants. We compared core DHS regions, which comprises a consensus centroid region aggregated across biosamples, and distances from the summit at 25bp, 50bp and 100bp up- and down-stream, with a median core length of 37bp (Supplemental Figure 4A, B). By using variants from the intersection with the GWAS catalogue benchmarked against molQTL variants, we calculated precision scores as fold change above GWAS catalogue alone. DHS cores have ∼1.5 fold precision, and this is not increased by extending to 100bp on either side of the summit (Figure 4A). Although a large number of footprints fall outside the DHS core, DNase footprints are found at a higher frequency nearer DHS summits (Figure 4B). Together, these data suggest a mechanism for functional variants related to the enrichment of DNA-binding motifs near DHS summits. When comparing centrality as a means to identify functional variants with DNase footprints and eRNA, the precision is even higher with the combined DNase footprints / eRNA markers (Figure 4A).

**Figure 4.**
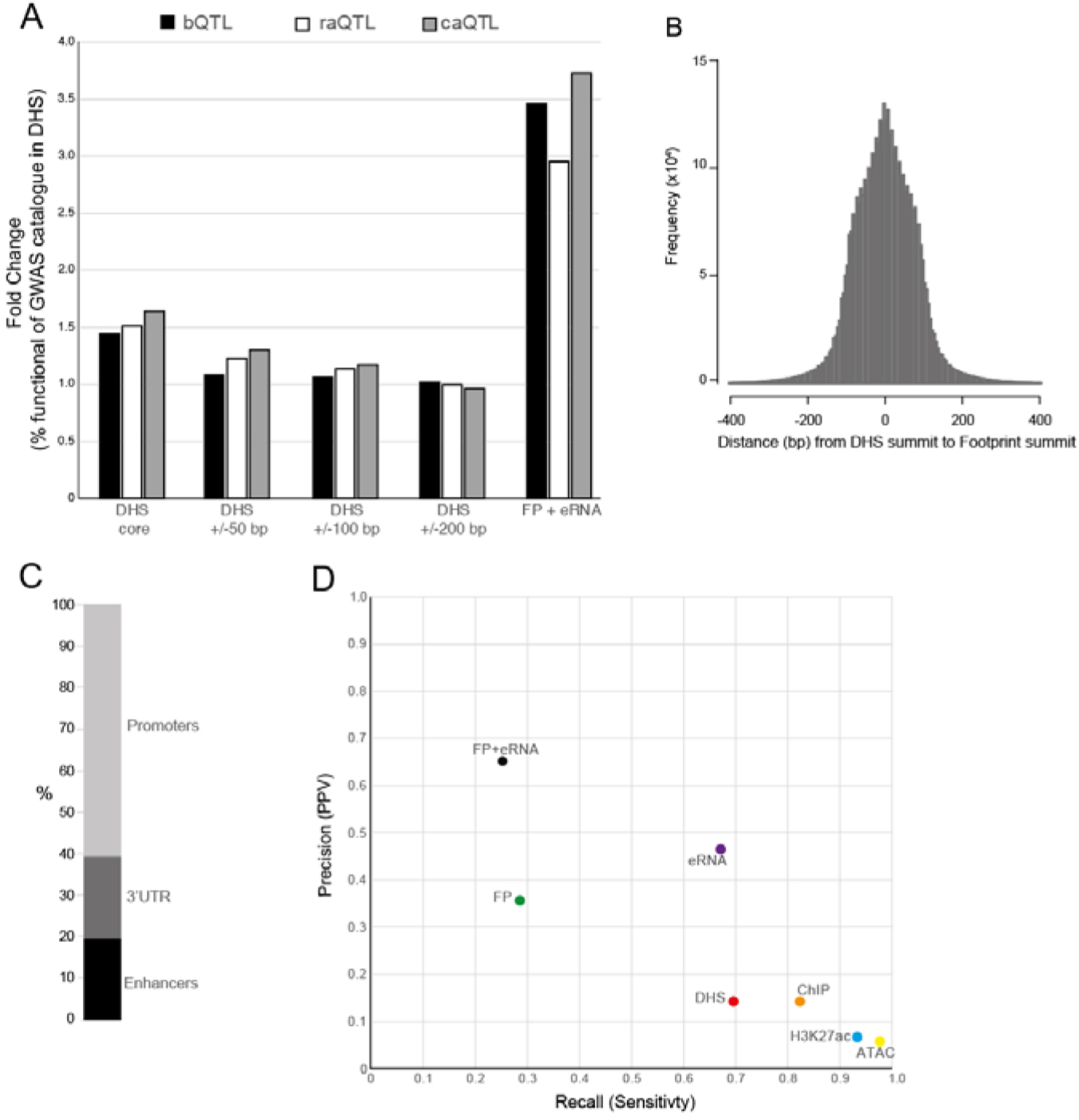
Centrality in the identification of functional variations, and benchmarking against non-coding Mendelian disease-associated variants. A. Precision score of DHS cores and DHS central lengths around summits, compared to DNase footprints with eRNA. Precision scores are expressed as fold change of precision score for the indicated feature over the precision score for GWAS variants alone. Scores are benchmarked for each molQTL. B. Distribution of nearest DHS footprint up- and downstream relative to DHS summits. C. Genomic annotation of validated, rare, non-coding Mendelian disease-associated variants. D. Precision-recall graph determined from 230 non-coding Mendelian disease-associated variants, with spike-in of random genomic variants from dbSNP (∼4K) to produce a validated variant probability of ∼0.05. Features were intersected to determine precision (positive predictive value – PPV) and recall (sensitivity). Binary outcomes were determined from feature intersections.

### Replication with functional, rare, non-coding Mendelian disease-associated variants

To validate our observation of high precision using the combination of DNase footprints and eRNA, we sought to replicate our analysis using a set of validated functional non-coding variants. We used a published set of manually curated rare non-coding variants associated with Mendelian diseases consisting of 210 non-redundant variants in promoter, intergenic, intronic or 3’UTR regions (Smedley et al., 2016) (Figure 4C). Given the small number of variants within this validated set, we limited our precision calculations to single markers, or the combination of DNase footprints and eRNA only. We determined precision by spike in of ∼4K randomly selected variants across the whole genome, giving a true positive rate of approximately random chance (∼0.05). We generated precision and recall scores based on the mean from 10 permutations of random genomic variant spike-ins (Figure 4D). The combination of DNase footprints and eRNA shows the highest precision at 0.65, but low recall (0.25). In contrast, ATAC-seq and H3K27ac show low precision but high recall. DNase footprints and eRNA each alone have intermediate precision and recall. These observations are consistent with our analysis of the GWAS catalogue and genomic variants, suggesting the utility of DHS footprints and eRNA in identifying both rare and common functional non-coding variants.

### Prioritising variants from genome-wide association studies

To demonstrate the utility of DNase footprints and eRNA in prioritising non-coding variants, we analyse 53 traits for intersections with footprints, eRNA, or both (Table 1). Across these traits the majority of associated variants are, as expected, in the non-coding genome, and this is maintained when using the combined footprint and eRNA marks to prioritise variants (Figure 5A). Using the leukocyte count trait from the GWAS catalogue to exemplify our prioritisation approach, ∼10K non-redundant variants with significance of < 9×10^-6^ from 51 studies is reduced ∼30 fold by requiring co-localisation with DNase footprints and eRNA (Table 1). Using gene ontology (GO) analysis of neighbouring genes, we applied GREAT analysis (Tanigawa et al., 2022) to all variants from the leukocyte count trait compared to just the variants within footprints and eRNA. GO terms are preserved between all variants and those in footprints and eRNA (Figure 5B), with enriched GO terms in keeping with leukocyte function. For the most significant GO term, regulation of immune response, we observe preservation of 55% of genes in the footprint and eRNA variant set, compared with all variants (Figure 5C) suggesting that, despite a ∼30 fold reduction in number of candidate variants, the enriched genes and loci, are largely preserved.

**Figure 5.**
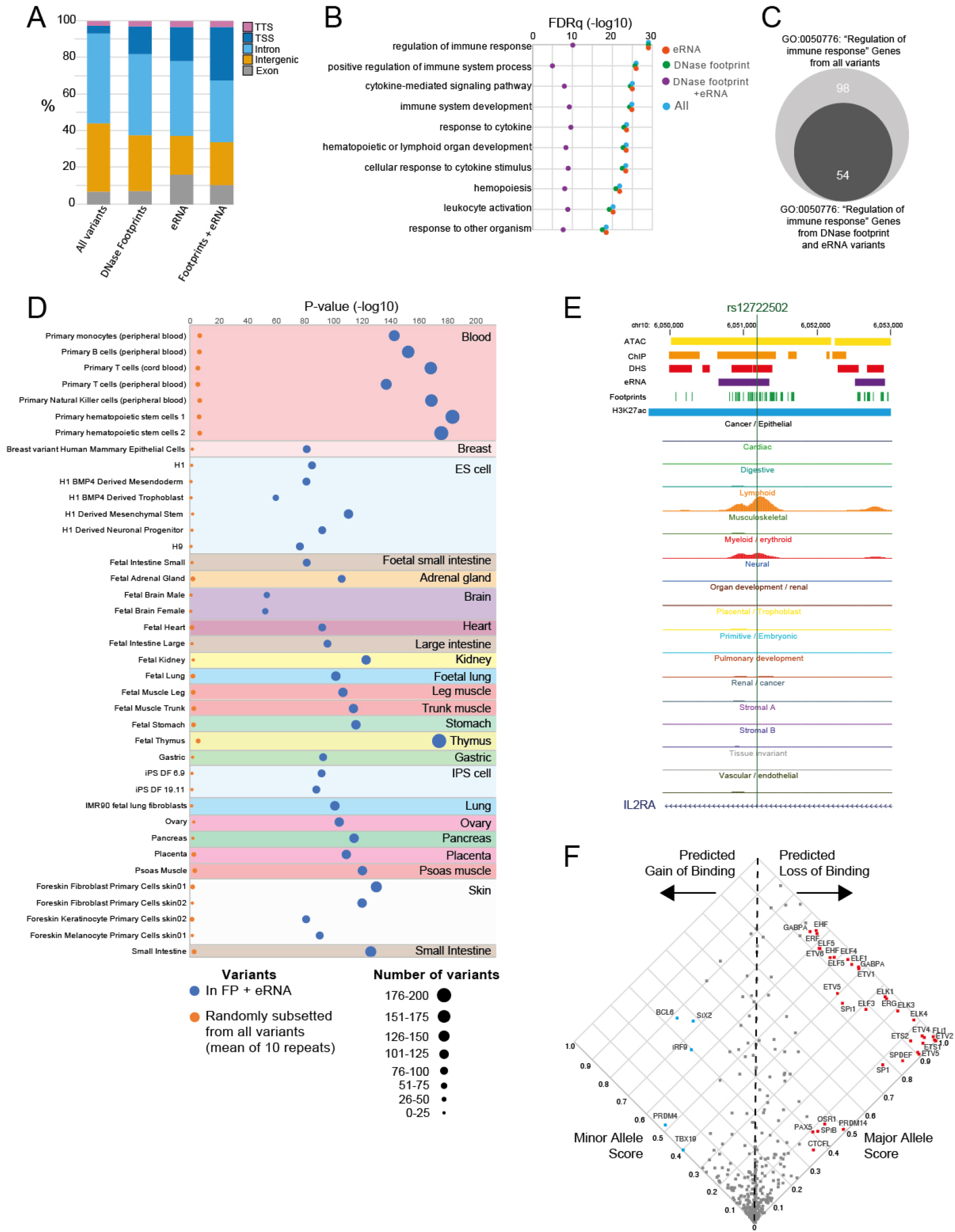
Prioritisation of variants using DNase footprints and eRNA preserve cell-specific features and functions. A. Genomic annotation of all variants for 53 traits from the GWAS catalogue, and annotations when intersection with DNase footprints, eRNA or both in combination. B. Gene ontology (GO) analysis of proximity genes relative to variants of lymphocyte count trait, for all variants for the trait from the GWAS catalogue, variants in DNase footprints, eRNA or both. Significance of the GO term is expressed as -log10 of the FDR q value (FDRq). Top ten GO terms for all variants are shown with FDRq for each intersection. C. Number of genes identified from GO analysis for the GO term “regulation of immune response” comparing number of genes from all variants compared to number of genes from variants in footprints and eRNA. D. Enrichment of cell-type specific active regulatory elements for the lymphocyte count trait, comparing variants in DNase footprints and eRNA, compared to an equal number (333) randomly selected variants from all lymphocyte count variants from the GWAS catalogue. The random selection was repeated ten times, and the mean P-value (-log10) shown. Enrichment analysis was determined using FORGE2. E. An example lymphocyte count-associated variant (rs12722502) which co-localises with a DNase footprint and eRNA. UCSC genome browser shot shows merged features, and ENCODE DNase-seq bigwig tracks for each cell-/tissue-type, for the IL2RA locus. F. Predicted impact of rs12722502 on DNA-binding factors. Scatter plot of the major and minor allele binding score for DNA-binding factor motifs showing predicted gain or loss of binding. Example DNA-binding factors are labelled.

**Table 1.** Co-localisation of GWAS traits from the GWAS catalogue with DNase footprints and eRNA. 53 GWAS traits from the GWAS catalogue, showing the number of studies, total number of variants, and number of non-redundant variants. Non-redundant variants were then intersected with DNase footprints, eRNA or both. The numbers co-localising are shown and the table sorted by non-redundant variant number.

| Trait | No of studies | Variants<br>Total | Variants<br>Non-redundant | Variants in FP | Variants in eRNA | Variants in FP/eRNA |
| --- | --- | --- | --- | --- | --- | --- |
| Leukocyte count (WBC) | 51 | 19264 | 10158 | 722 | 1501 | 333 |
| Body Height (exc Child traits) | 53 | 7487 | 6038 | 341 | 653 | 141 |
| Body Mass Index (BMI) (exc Child traits) | 114 | 8672 | 4922 | 195 | 344 | 56 |
| HDL Cholesterol (HDL) | 123 | 10872 | 3969 | 200 | 503 | 78 |
| Bone density (BD) | 63 | 5883 | 3755 | 190 | 351 | 63 |
| Triglycerides (TRIG) | 114 | 8567 | 3737 | 184 | 423 | 72 |
| Type 2 diabetes (T2DM) | 133 | 5679 | 3612 | 172 | 326 | 59 |
| Systolic blood pressure (SBP) | 73 | 4227 | 3096 | 173 | 341 | 73 |
| FEV1:FVC ratio | 23 | 3283 | 2957 | 121 | 165 | 36 |
| Mean corpuscular haemoglobin (MCV) | 29 | 4206 | 2779 | 201 | 508 | 121 |
| Platelet count (PLT) | 28 | 3686 | 2545 | 187 | 387 | 89 |
| Glomerular filtration rate (GFR) | 30 | 3726 | 2442 | 159 | 287 | 68 |
| Alzheimer's disease (AD) | 88 | 2811 | 2380 | 115 | 231 | 1 |
| Monocyte count (MC) | 20 | 3474 | 2300 | 179 | 375 | 83 |
| Asthma | 93 | 3442 | 2275 | 116 | 286 | 65 |
| Diastolic blood pressure (DBP) | 69 | 3031 | 2203 | 106 | 223 | 43 |
| Coronary artery disease (CAD) | 72 | 3037 | 2062 | 119 | 203 | 47 |
| Breast carcinoma (BC) | 100 | 2566 | 1852 | 66 | 163 | 20 |
| Mean platelet volume (MPV) | 17 | 2046 | 1423 | 104 | 283 | 64 |
| Inflammatory bowel disease (IBD) | 78 | 2075 | 1134 | 58 | 148 | 29 |
| Rheumatoid arthritis (RA) | 69 | 1760 | 1094 | 70 | 160 | 40 |
| Prostate carcinoma (PC) | 57 | 1577 | 948 | 61 | 107 | 31 |
| Systemic lupus erythromatosis (SLE) | 41 | 1197 | 936 | 49 | 145 | 27 |
| Vital Capacity (VC) | 17 | 916 | 844 | 0 | 0 | 0 |
| Lung carcinoma (LC) | 55 | 1058 | 823 | 24 | 73 | 15 |
| Crohn's Disease (CD) | 45 | 995 | 747 | 43 | 96 | 20 |
| Multiple sclerosis (MS) | 41 | 819 | 727 | 48 | 123 | 23 |
| Psoriasis | 28 | 895 | 691 | 44 | 98 | 20 |
| Chronic Kidney Disease (CKD) | 40 | 1111 | 689 | 33 | 63 | 13 |
| Type 1 diabetes (T1DM) | 41 | 815 | 658 | 28 | 83 | 18 |
| Heart rate (HR) | 27 | 792 | 582 | 22 | 41 | 10 |
| Age at menarche (Men) | 16 | 636 | 579 | 27 | 35 | 6 |
| Cholesterol:total lipids ratio (Chol) (exc Child traits) | 2 | 1054 | 568 | 36 | 89 | 14 |
| Ulcerative colitis (UC) | 28 | 753 | 556 | 33 | 84 | 17 |
| Atrial fibrillation (AF) | 20 | 767 | 477 | 32 | 50 | 11 |
| Stroke | 30 | 618 | 477 | 19 | 31 | 4 |
| Ovarian carcinoma (OC) | 19 | 600 | 448 | 19 | 24 | 5 |
| Parkinson's Disease (PD) | 50 | 546 | 434 | 19 | 42 | 7 |
| Melanoma | 26 | 519 | 392 | 23 | 33 | 8 |
| Eczema | 4 | 505 | 391 | 28 | 69 | 16 |
| Ankylosing spondylitis (AS) | 9 | 416 | 370 | 25 | 50 | 13 |
| Primary sclerosing cholangitis (PSC) | 6 | 299 | 282 | 26 | 43 | 15 |
| Hypothyroidism | 7 | 292 | 259 | 17 | 35 | 7 |
| Age at menopause (AAM) | 11 | 335 | 254 | 12 | 40 | 5 |
| Amyotrophic lateral sclerosis (ALS) | 27 | 262 | 223 | 12 | 13 | 3 |
| Celiac Disease (CEL) | 13 | 249 | 218 | 10 | 27 | 3 |
| Primary biliary cirrhosis (PBC) | 8 | 252 | 196 | 12 | 39 | 5 |
| Juvenile idiopathic arthritis (JIA) | 11 | 148 | 145 | 3 | 15 | 1 |
| Colorectal carcinoma (CC) | 7 | 129 | 127 | 8 | 12 | 3 |
| Idiopathic pulmonary fibrosis (IPF) | 6 | 70 | 58 | 7 | 16 | 4 |
| Glioma | 9 | 153 | 54 | 4 | 8 | 2 |
| Renal carcinoma (RC) | 7 | 50 | 39 | 2 | 4 | 0 |
| Narcolepsy (NAR) | 8 | 36 | 31 | 1 | 5 | 0 |

Non-coding variants function in a cell-type specific manner (e.g. Rai et al., 2020; Torres et al., 2020) but uncovering the affected cell-type from GWAS can be challenging. By leveraging DNase footprints and eRNA to identify functional variants with high precision, we can then infer the relevant cell-type from this subset of variants. As both footprints and eRNA are associated with open chromatin (Young et al., 2017), we took the leukocyte count associated variants co-localising with footprints and eRNA and identified cell-type specific enrichment of DHS using FORGE2 (Breeze et al., 2022). Since footprints are derived from DNase-seq, the subset of variants that colocalise with DNase footprints and eRNA show a greater enrichment in DHS compared to an equal numbers of randomly selected GWAS variants for leukocyte count. Importantly however, despite our cell- and tissue-type agnostic approach of merging features, colocalised variants show enrichment in haematopoietic or immune cell types (Figure 5D), demonstrating preservation of the ability to identify relevant cell-types associated with the trait. Highlighting one variant identified from DNase footprints and eRNA, rs12722502 is an intronic variant of *IL2RA*, and is in a myeloid and lymphoid specific DHS (Figure 5E). This variant is predicted to cause a loss of ETS family TF binding (Figure 5F), members of which are known to regulate immune cell function (Garrett-Sinha, 2013). Indeed, rs12722502 is associated with five-fold reduction in binding of ELF1 (a TF in the ETS family), in a human leukemic T-cell (Schwartz et al., 2017).

**Figure 6.**
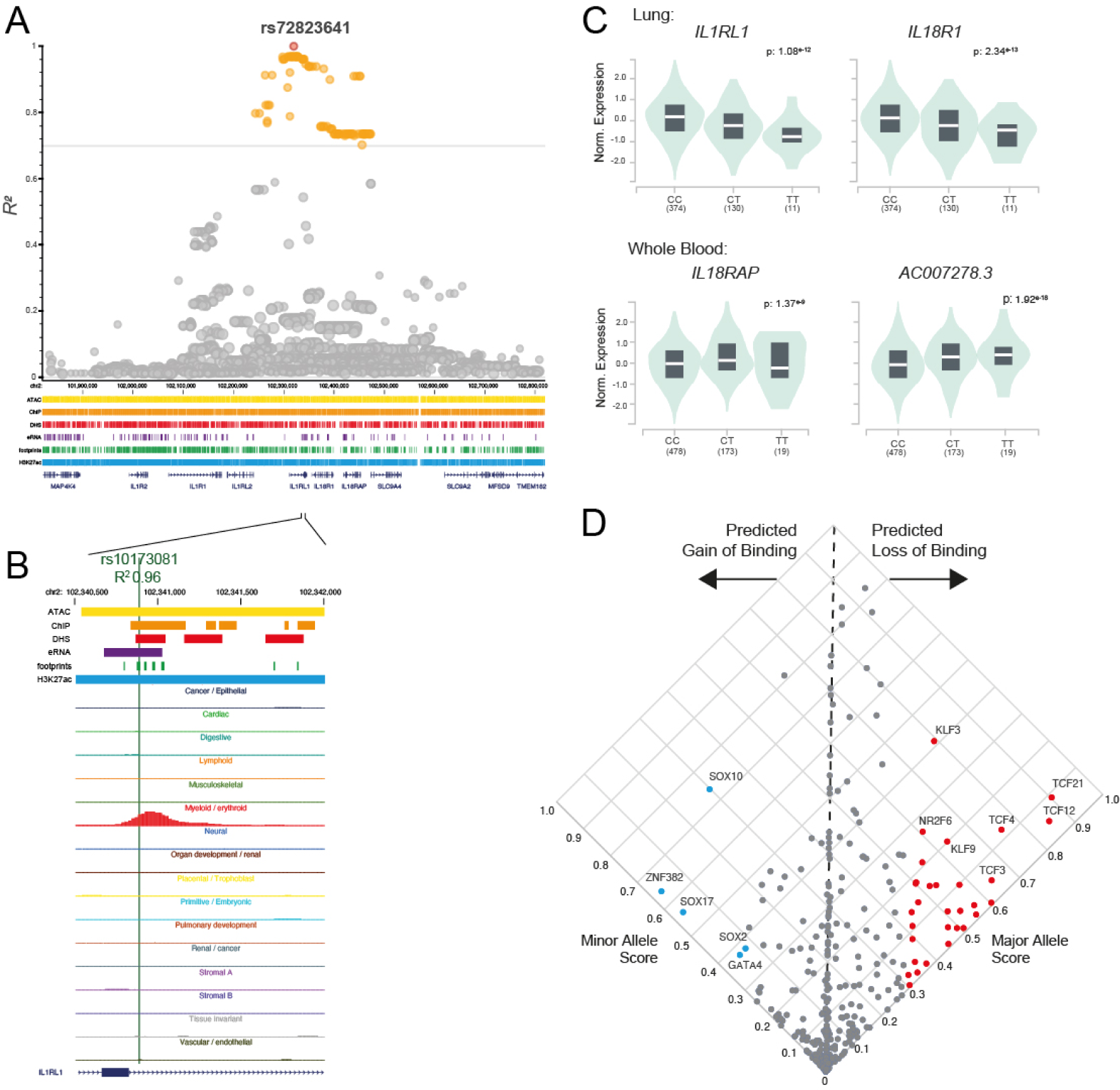
DNase footprints and eRNA can prioritise functional variants for variants in LD with a lead variant. A. Manhattan plot of the asthma-associated rs72823641 variant (red), with variants in LD >= 0.7 indicated in orange. Merged feature tracks are shown for the locus. Genome co-ordinates (Mb) are indicated. B. rs10173081 overlaps a DNase footprint and eRNA. UCSC genome browser shot for the intronic region of IL1RL1 with rs10173081 overlapping a myeloid specific DNase hypersensitive site. C. rs10173081 is an expression QTL (eQTL) for multiple genes in lung and whole blood. GTex violin plots for normalise expression of the indicated genes across homozygous major allele, heterozygous major and minor allele, and homozygous minor allele. D. Predicted impact of rs10173081 on DNA-binding factors. Scatter plot of the major and minor allele binding score for DNA-binding factor motifs showing predicted gain or loss of binding. Example DNA-binding factors are labelled.

### Identifying functional variants in linkage disequilibrium with lead variants

Uncovering functional or causative variants from GWAS is also challenged by LD, whereby the causative variant may be a variant in LD with the GWAS lead variant. To investigate the utility of colocalization with DNase footprints and eRNA to prioritise variants in LD with lead variants, we selected the asthma trait, and considered strong lead GWAS variants which were not associated with the *HLA* locus, to avoid cell-type bias. We identified rs72823641, which has been found to be associated with asthma in multiple GWAS (Johansson et al., 2019), with p-values ranging from 4 x 10^-12^ to 2 x 10^-136^, and mapping to a large intron of *IL1RL1*. We found 92 variants in LD with rs72823641 (R^2^ > 0.7) using European ancestry data (Figure 6A). Intersecting these with DNase footprints and eRNA reduced the variants to one candidate - rs10173081 (R^2^ = 0.96), which resides in myeloid specific open chromatin (Figure 6B). While rs10173081 is not tagged in the GWAS catalogue, it has been observed in other GWAS for asthma (Portelli et al., 2020). rs10173081 is also an eQTL for *IL1RL1* and *IL18R1* in lung, but also for *IL18RAP* and a non-coding RNA AC007278.3 in whole blood (Figure 6C). Applying TF motif based binding prediction, we identified putative loss of binding for TFs including TCF21, TCF12, TCF4 and KLF3 (Figure 6D). Interestingly, *TCF21* and *KLF3* loci have been previously identified as asthma risk-associated GWAS loci (Daya et al., 2019). In conclusion, by using combined DNase footprint and eRNA markers, plausible functional variants can be prioritised from variants sets in LD with lead variants.

## Discussion

GWAS have been instrumental in uncovering complex trait- and disease-associated variants and regulatory pathways. However, the identification of functional and causative variants remains a significant challenge, impacting the mechanistic understanding of complex disease genetics, and the identification of disease/trait relevant pharmacological targets. Few GWAS have been translated into therapeutic interventions (Trajanoska et al., 2023). As most variants associated with complex traits are in the non-coding genome, discovering functional variants is constrained by methodological limitations. Computational approaches have been developed that utilise variable genomic features to aid variant prioritisation, including variable use of functional annotations, conservation scores, and TF motifs (Wang et al., 2022, Dong et al., 2023). However, for non-coding variants these methods tend to have low precision for identifying functional variants (Wang et al., 2022).

Here, we systematically analysed six assays for active regulatory elements, benchmarking against independent molQTLs for enhancer activity. We find that ATAC peaks and H3K27ac are poor predictors. This is in keeping with previous studies that have shown that deletion of elements harbouring candidate variants marked by ATAC peaks or H3K27ac are unable to detect functional impact (Gasperini et al., 2017, Downes et al, 2021, Chen et al., 2023). We hypothesise that the poor precision but high recall of ATAC-seq and H3K27ac peaks relates to their high feature size and high genomic coverage. In contrast, we identify a signature that performs well with respect to precision in identifying genetic variants with putative function. Individually DNase footprints and eRNA both show good precision, however their combination further improves precision. DNase footprints, and eRNA have independently been suggested to be enriched for putative functional variants (Vierstra et al., 2020; Yao et al., 2022). However, we demonstrate that their combination provides superior discrimination.

Our model for the precision offered by combined DNase footprints and eRNA markers is based on direct DNA binding of a factor, such as a TF, and that such binding recruits RNA polymerase and other machinery to promote regulatory activity. In agreement with this model, an analysis of ATAC peaks in primary CD4+ T-cells showed <30% demonstrated activity using an MPRA, with MPRA signals correlating well with eRNA production (Stefan and Barksi, 2023), demonstrating an abundance of inactive accessible regulatory elements. Additionally, MPRA of glucocorticoid receptor binding sites showed <20% had significant activity (Vockley et al., 2016), suggesting that many bound TFs are not productive. If evidence of binding is required for functional variant prediction, then ChIP-seq signals would be expected to perform well. However, artefactual binding observed from cross-linking in ChIP-seq experiments (Mercer et al., 2013), and co-localisation of multiple TF binding events enriching for non-related motifs, can lead to false positive binding sites (Partridge et al., 2020). Therefore, the footprint signal of direct TF binding, with a detection of eRNAs, provides evidence for an active enhancer. Variants within these regions, particularly in DNA sequences interfacing with TF DNA binding domains (DBD), are thus more likely to be functional.

Despite the high precision for functional variants using the combination of DNase footprint and eRNA, restricting to these two markers comes with limitations, principally related to low recall, causing a high number of false negatives. The low recall can be explained by limitations inherent to the DNase footprint and eRNA datasets. Firstly, the identification of DNase footprints may be related to residency time of TFs on DNA, where rapidly exchanging factors impart poor footprints (Sung et al., 2014). Variants associated with altering binding of dynamic factors may therefore be missed. To overcome this, detection of footprints could be improved by enzymatic digestion bias correction (Calviello et al., 2019). Secondly, the eRNA database used in our analysis represents the lowest number of biosources, and is therefore incomplete, with highly variable numbers of features between cell-types, and predominantly under basal conditions, which would exclude context-dependent enhancer activity (Yao et al., 2022, Vockley et al., 2016). To address this, eRNA profiling, for example using PRO-seq, is required in cell-types, and under conditions, relevant to the GWAS trait of interest.

Together, we present our findings as a framework, FINDER - Functional SNV IdeNtification using DNase footprints and Enhancer RNA (Figure 7). In this framework, the combination of DNase footprints and eRNA can prioritise and reduce candidate variants to a set with high precision. The merged datasets used can be analysed for relevant cell-types associated with the complex trait or disease. This has important implications towards current computational approaches, and databases for predictive functional variant scores. Our findings can inform predictive scoring approaches as these do not utilise eRNA as a predictive marker. Additionally, predictive algorithms may use summative scoring for feature overlaps, whereas our analysis suggests that specific combination of markers may be non-linear in their prediction of variant function. Finally, the high precision of DNase footprints and eRNA compared to other marks may be used for functionally informed fine mapping approaches (Kumasaka et al., 2016), which could improve detection of functional, causative non-coding variants.

**Figure 7.**
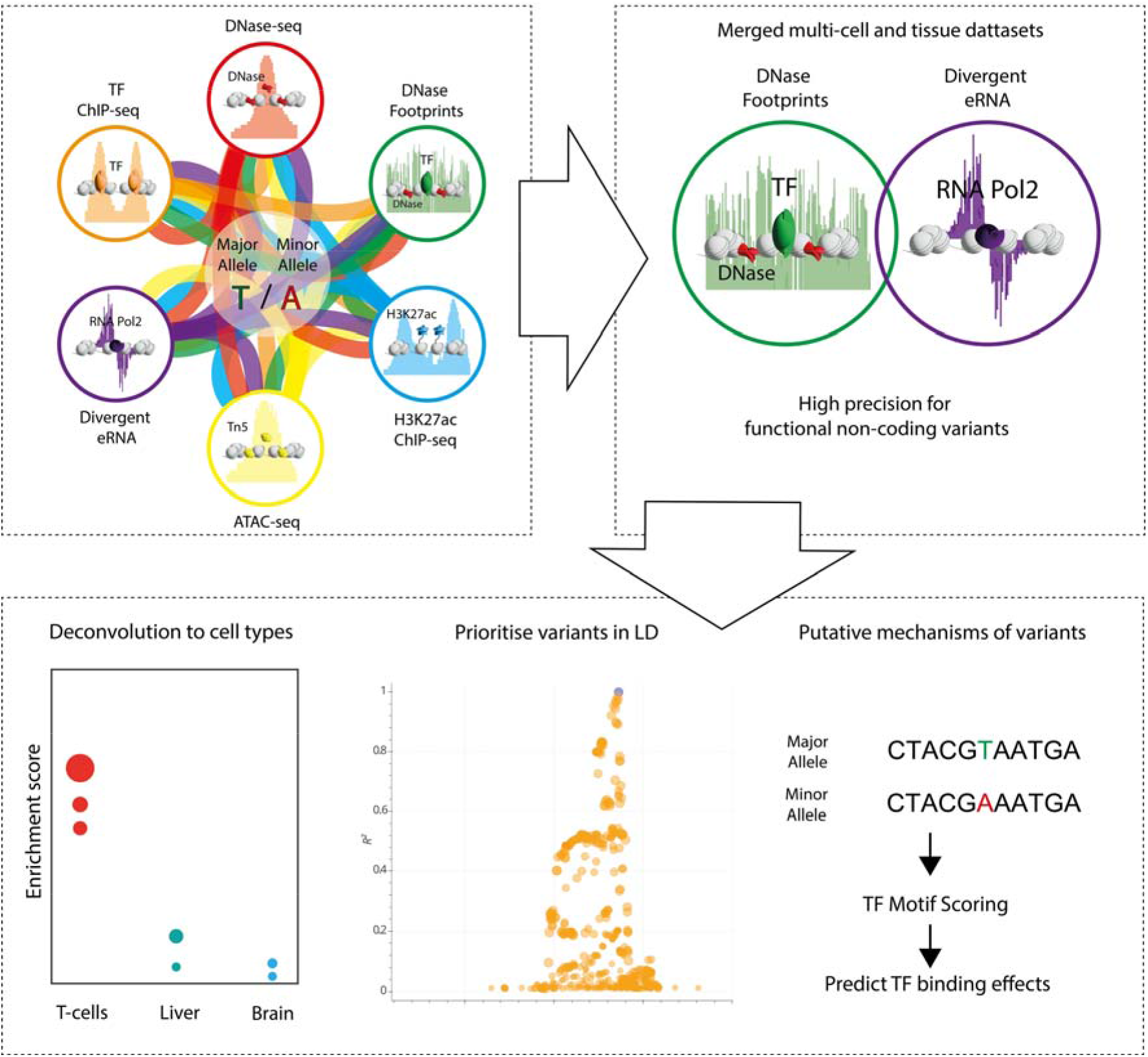
The FINDER framework - Functional SNV IdeNtification using DNase footprints and Enhancer RNA. Markers of active regulatory elements as merged datasets from multiple cell- and tissue-types are used to predict functional variants. The combination of DNase footprints and eRNA provide the highest precision, which can be applied to GWAS traits or to predict functional variants in linkage disequilibrium (LD). Predicted variants can then be interrogated for cellular and molecular function, by deconvolving marker enrichment to predict relevant cell-types, or the predict altered binding affinity for DNA-binding factors.

## Methods

### Data sources

Datasets for active chromatin marks were obtained from published and publicly available sources. DNase hypersensitive sites (Meuleman et al. 2020) and DNase footprints (Vierstra et al., 2020) were obtained from ENCODE. H3K27ac was obtained from GTRD (Kolmykov et al., 2021) and dbInDel (Huang et al., 2020). Human ChIP-seq and ATAC-seq datasets were obtained from GTRD (Kolmykov et al., 2021). Enhancer RNAs were obtained from the PINTS portal (Yao et al., 2022). These data sources provide uniformly analysed datasets for downstream analysis. Details of downloaded versions and sources are shown in Supplementary table 1. For each type of active chromatin mark, to generate assay master files agnostic to multiple cell-types or factor in the case of ChIP, Bedtools merge version from the bedtools suite v2.30.0 (Quinlan and Hall, 2010) was used, requiring overlap of 1 bp or greater.

QTL datasets were obtained from published datasets. Binding QTL (bQTL) were downloaded from AD ASTRA (Abramov et al., 2021) which determines allele-specific binding events from human ChIP-seq datasets and identifies genetic variants from ChIP-seq data. Reporter assay QTL (raQTL) data were downloaded from publicly available massively parallel reporter assays (MPRA) (van Arensberggen et al., 2019). Chromatin accessibility QTL (caQTL) were downloaded from ENCODE (Vierstra et al., 2020) and QTLbase (Zheng et al., 2020). Details of downloaded versions and sources are shown in Supplementary table 1. For each molecular QTL, where an rsID occurs in multiple datasets, either multiple assays (such as ChIP) and multiple cell types, these were collapsed to generate a list of unique rsIDs.

GWAS summary statistics were downloaded from the NHGRI-EBI GWAS catalog V1.0.2 (Sollis et al., 2023), using variants with a p-value <10^-5^. Where analysis presented is agnostic to traits or studies, a master list of rsID found in the GWAS catalog was generated by collapsing to a list of unique rsIDs. Details of downloaded version and source are shown in supplementary table 1.

The dataset for genetic variants across the human genome was obtained from UCSC (Sherry et al., 2001) using the dbSNP build 151 containing uniquely mapping common SNPs with a >= 1% minor allele frequency (MAF). SNPs were randomly subsetted using gshuf in command line.

### Confusion matrix calculations and normalisations

Precision scores were calculated as the fraction of the number of variants (GWAS catalogue or other variant sets as indicated) overlapping the benchmarked molQTL (True Positives) over the total number of variants (True and False Positives) intersecting the feature (or feature combinations). Precision scores were expressed as a Z-score or fold change. Clustering of precision z-scores was performed in R using heatmap cluster function with the Ward D2 method. Fold changes were expressed as the precision score of the candidate set intersected with features divided by precision score of the candidate set without intersection with features.

### Genome build conversions

Data were analysed using human genome build Hg38. Where primary data sources were aligned to other genome versions (Hg 19), data was converted to Hg38 using Liftover (Hinrichs et al., 2006). The genome version of the primary data sources are indicated in supplementary table 1.

### Intersections of features

To analyse the overlap between genetic variants, and/or genomic features, Bedtools intersect. from the bedtools suite v2.30.0 (Quinlan and Hall, 2010) was used. For intersections with rsID, these were considered as the first feature (-a) in bedtools intersect - a <bed/vcf > -b <bed>.

### Random forest classification

The PeakPredict package (https://github.com/efriman/PeakPredict) command overlap_peaks was run with settings ‘--predict_column molQTL --model RandomForestClassifier --balance --shap’, where balancing downsamples the groups to have the same size prior to splitting into test and training sets. The PeakPredict package implements scikit-learn (Pedregosa et al., 2011) and SHAP values.

### Gene Ontology analysis

GO analysis utilised Genomic Regions Enrichment of Annotations Tool (GREAT) version 4.0 (Tanigawa et al., 2022) with rsID converted to BED format, using the default settings.

### Predicting Functional base on Motif

Predictions of TF binding based on motifs utilised the FABIAN-variant (Steinhaus et al., 2022) web interface, using Hg38m, using “All” TFS, the “TFFM detailed” model, and the JASPAR2022 database.

### Linkage disequilibrium analysis

To generate variants in LD with a lead variant, we used LDproxy with the GRCh38 genome build, using R^2^ to identify LD, and pan-European ancestry. Variants with R^2^ >= 0.7 were considered for further downstream analysis.

### Cell-type specific analysis

To identify cell-type enrichment of variants, we used FORGE2 (Breeze et al., 2022), with input as rsID of variants, using DHS as data for enrichement, without LD filtering of rsIDs.

### Genome location mapping

To annotate the genomic features to genome location, we used the annotatePeaks.pI program from HOMER tools v4.1 (Heinz et al., 2010). Gene annotation was customised using a GTF file containing hg38 gene transcript sets, downloaded from UCSC (hgdownload.soe.ucsc.edu/goldenPath/hg38/bigZips/genes/hg38.refGene.gtf.gz).

## Data Availability

Individual data sets are described in “Data sources” of the Methods section. The merged datasets for each active regulatory marker, merged as described in “Data sources”, can be found at https://github.com/sbiddie/FINDER.

## Author contributions

SCB and WAB conceived the study. SCB, GW, EFH and ETF performed data analyses. SCB wrote the manuscript. All authors were involved in the preparation of the manuscript.

**Supplemental figure 1.**
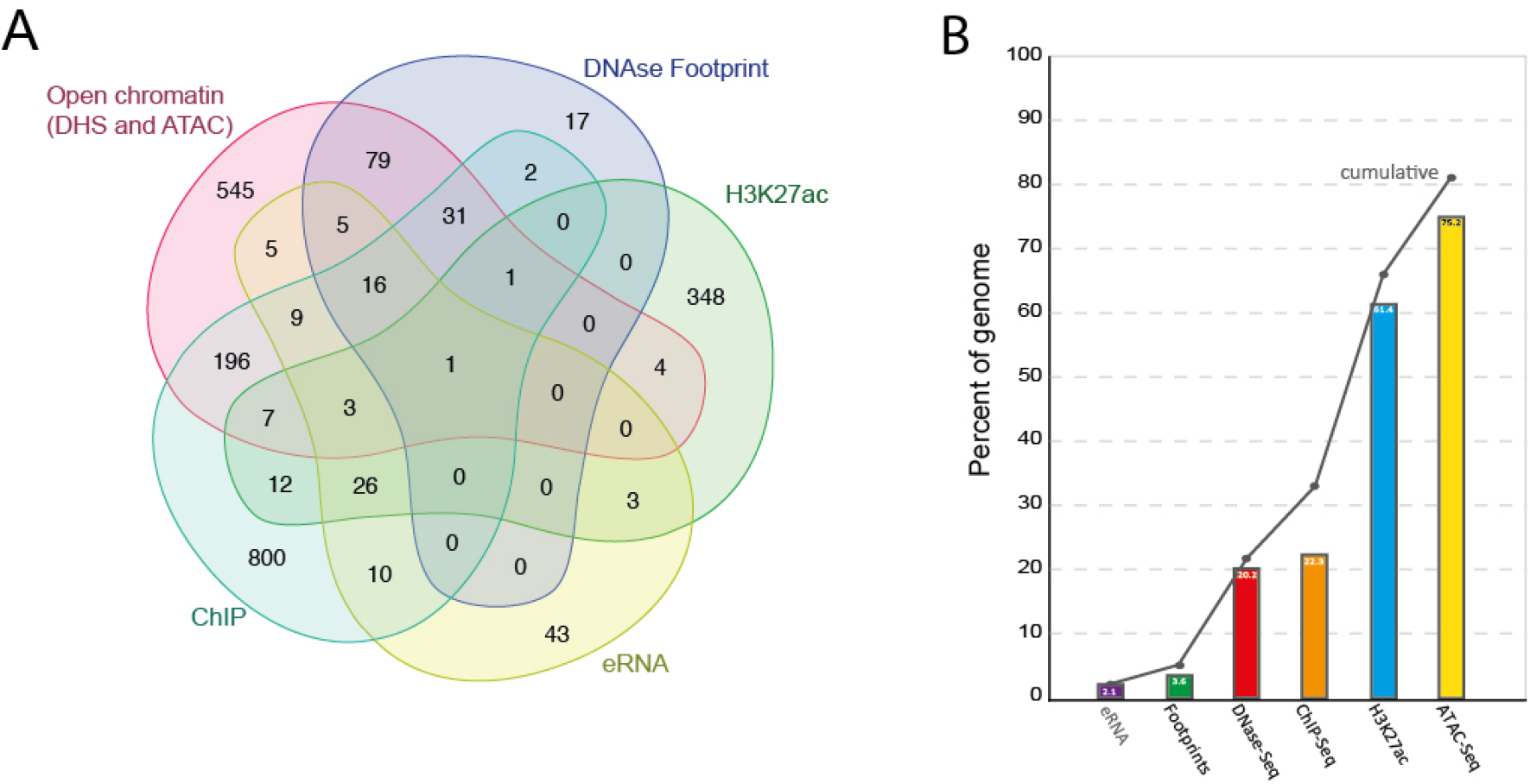
Features of markers of active regulatory elements. A. Euler diagram for intersections of biosources for each type of feature. DNase-seq (DHS) and ATAC-seq have been combined for visualisation. B. Percentage of genome coverage of total number of genomic basepairs (Hg38 build), with the percentage for each feature and the cumulative percentage shown.

**Supplemental figure 2.**
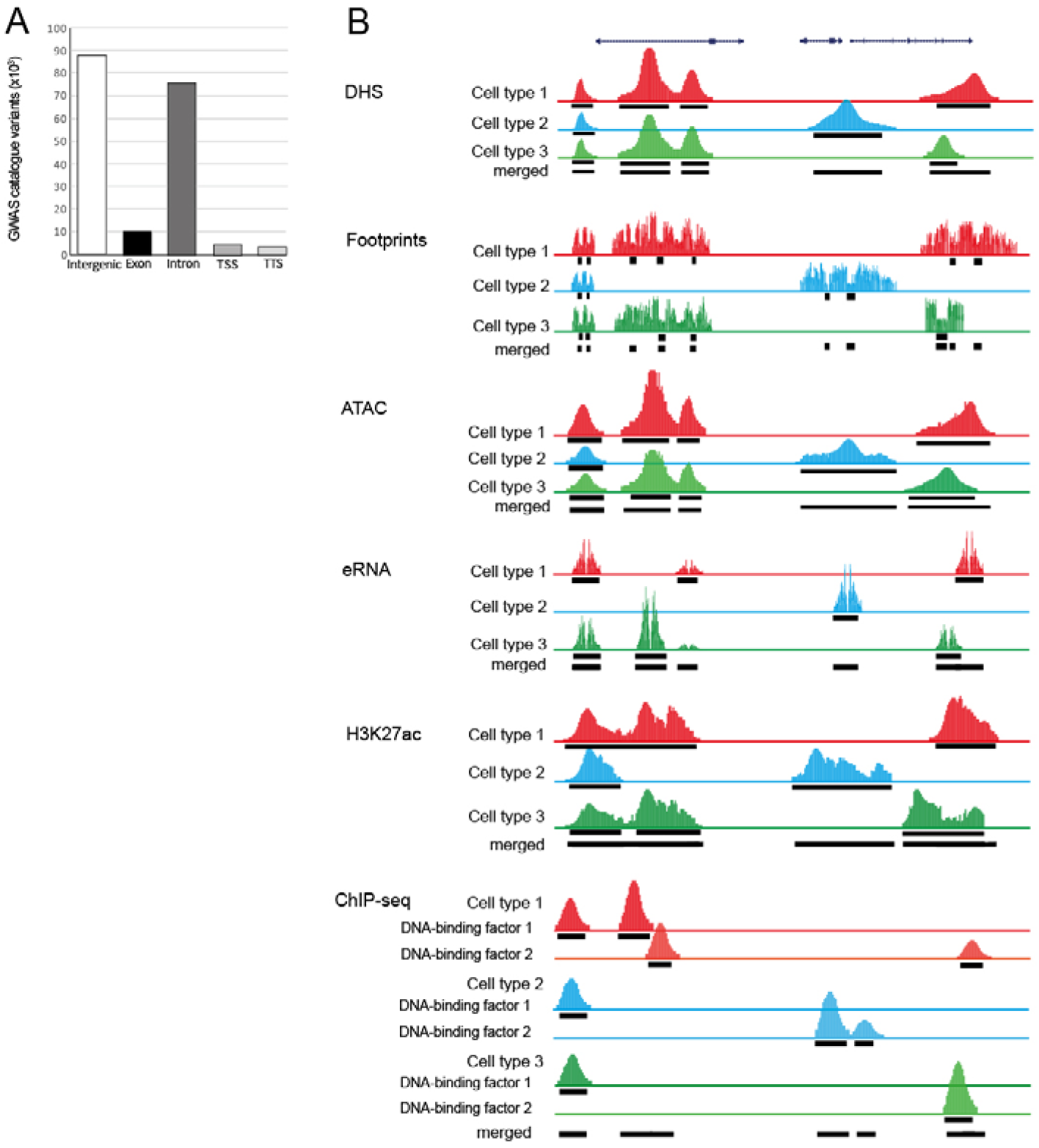
Genomic features of the GWAS catalogue and generation of merged active regulatory element datasets. A. Genomic annotation of variants from the GWAS catalogue showing that the majority reside in the non-coding genome. B. Illustrated examples of merging of different biosources for the same feature to generate a cell- and tissue-agnostic set.

**Supplemental figure 3.**
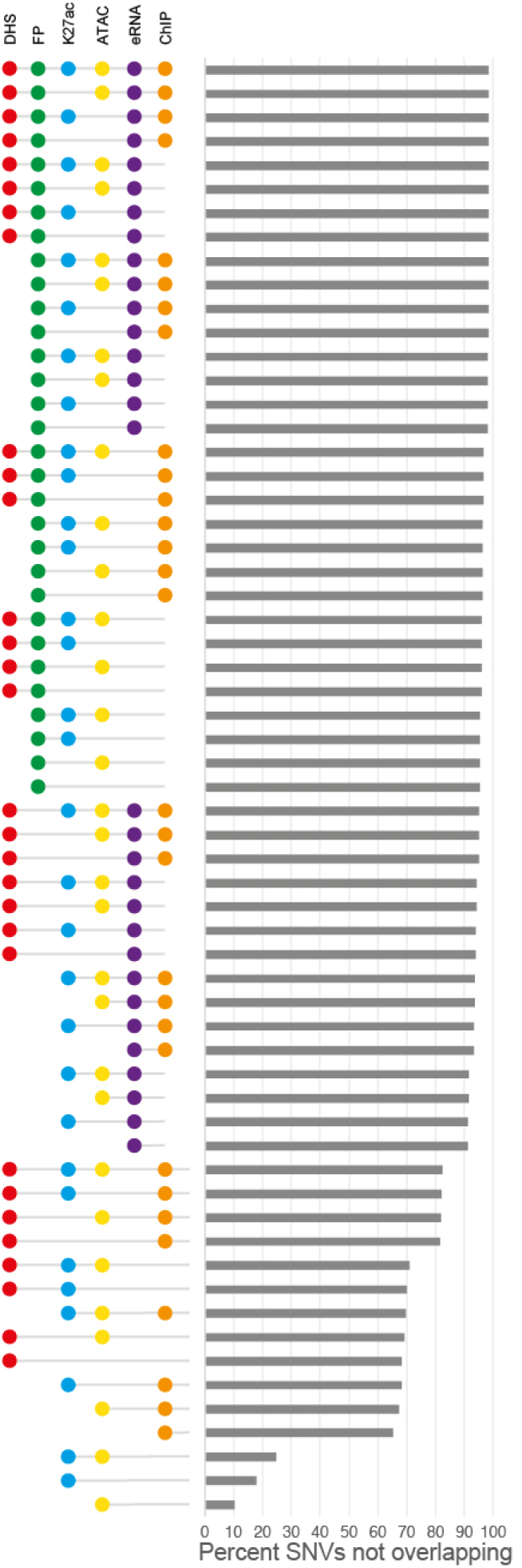
Percentage of the GWAS catalogue (∼181K) variants for each features, and combinations, classed as true negatives.

**Supplemental figure 4.**
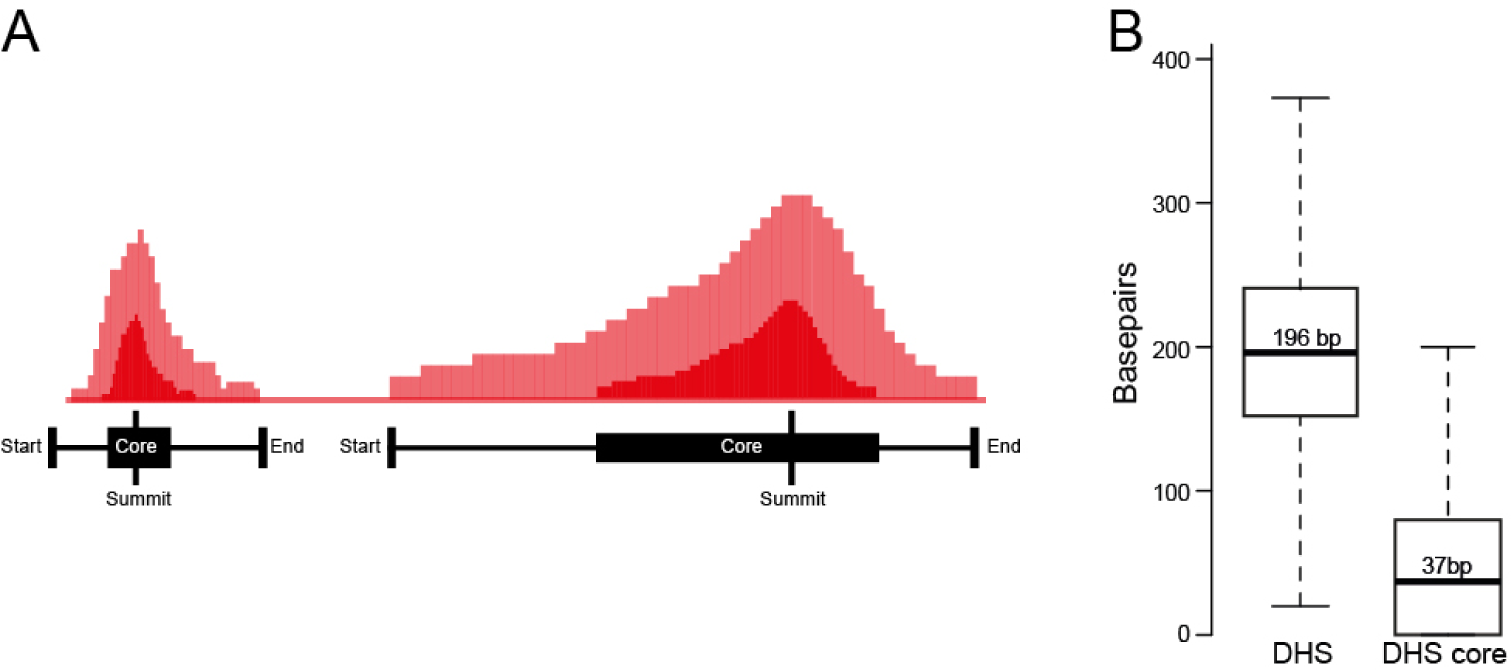
Features of DHS centrality measures. A. Schematic of DNase hotspots showing summits, DHS cores, and start and end locations. B. Boxplot of unmerged DHS lengths compared to DHS cores.

**Supplementary table 1. Data sources.** The table shows the source, version and genome build for each dataset. Where the genome version differs, data has been converted to Hg38 using Liftover (see methods).

